# Three-dimensional traction microscopy accounting for cell-induced matrix degradation

**DOI:** 10.1101/842831

**Authors:** Dawei Song, D. Thomas Seidl, Assad A Oberai

## Abstract

Tractions exerted by cells on the extracellular matrix (ECM) are critical in many important physiological and pathological processes such as embryonic morphogenesis, wound healing, and cancer metastasis. Three-dimensional Traction Microscopy (3DTM) is a tool to quantify cellular tractions by first measuring the displacement field in the ECM in response to these tractions, and then using this measurement to infer tractions. Most applications of 3DTM have assumed that the ECM has spatially-uniform mechanical properties, but cells secrete enzymes that can locally degrade the ECM. In this work, a novel computational method is developed to quantify both cellular tractions and ECM degradation. In particular, the ECM is modeled as a hyperelastic, Neo-Hookean solid, whose material parameters are corrupted by a single degradation parameter. The feasibility of determining both the traction and the degradation parameter is first demonstrated by showing the existence and uniqueness of the solution. An inverse problem is then formulated to determine the nodal values of the traction vector and the degradation parameter, with the objective of minimizing the difference between a predicted and measured displacement field, under the constraint that the predicted displacement field satisfies the equation of equilibrium. The inverse problem is solved by means of a gradient-based optimization approach, and the gradient is computed efficiently using appropriately derived adjoint fields. The computational method is validated *in-silico* using a geometrically accurate neuronal cell model and synthetic traction and degradation fields. It is found that the method accurately recovers both the traction and degradation fields. Moreover, it is found that neglecting ECM degradation can yield significant errors in traction measurements. Our method can extend the range of applicability of 3DTM.

## 1. Introduction

Tractions exerted by cells not only guide mechanical events such as the folding, stretching, and repositioning required for tissue morphogenesis [1, 2], but also activate signaling pathways that regulate basic cellular processes such as survival, proliferation, and stem-cell differentiation [3, 4, 5, 6, 7, 8]. Therefore, precise measurement of cellular tractions is crucial for understanding the role of mechanical forces in various physiological and pathological processes [9, 10]. For instance, the magnitude and direction of tractions have been found to modulate translation and transcription of vascular endothelial cells [11]; the temporal evolution of tractions have been found to control the speed and direction of a migrating cell [12, 13].

Over the years, several methods have been developed to estimate tractions imparted by cells. Interested readers are referred to several comprehensive review articles on traction measurements [9, 10, 14, 15, 16, 17]. Among various techniques, three-dimensional traction microscopy ^1^ (3DTM) has the distinguishing feature that it can be used to quantify cellular tractions in 3D settings that are similar to the *in vivo* cell environment [18]. In 3DTM, cells are fully encapsulated in the extra-cellular matrix (ECM), which contains fluorescent micro-beads. After the cell has spreaded into the matrix and applied tractions, a set of microscopy images are generated to measure the morphology of the cell and the locations of the beads. Tractions are then fully removed by chemically lysing the cell, causing the matrix to relax to its undeformed state. Another set of images are generated to track the locations of the beads in the relaxed state of the matrix. Making use of the two sets of images acquired before and after cell lysis, displacements of the beads in response to cellular tractions are determined by using image cross-correlation techniques. Given the measured bead displacements, as well as a constitutive model that characterizes the mechanical response of the ECM, tractions are determined at the cell-ECM interface.

Most 3DTM approaches have assumed that the ECM has spatially uniform mechanical properties, which are typically measured by standard techniques such as bulk rheometry [18], indentation testing [19], and atomic force microscopy [20] prior to the encapsulation of the cell. However, since cells express enzymes (e.g., matrix metalloproteinase (MMP)) that can locally degrade the ECM [21, 22, 23, 24, 25], the assumption of uniform mechanical properties is questionable and is likely to introduce inaccuracies in estimating tractions. In particular, experimental studies have highlighted the importance of incorporating ECM heterogeneity in measuring stresses within the ECM [26, 27]. On the other hand, since the cell-induced changes are concentrated near the cells [21], it is challenging to directly measure the changes in mechanical properties using the techniques discussed above. Motivated by this, in this work we develop a computational approach that is able to accurately recover cellular tractions, as well as the mechanical heterogeneities in the ECM induced by cell remodeling. Remarkably, our approach does not require any additional data than what is collected in a typical 3DTM experiment.

Our approach is based on an inverse problem formulation [18, 28, 29, 30, 31], where the objective is to determine the traction field and a spatial distribution of mechanical properties that produces a displacement field that best matches the measured displacements. We first review relevant literature in inverse problems for 3DTM, keeping in mind that most studies assume a homogeneous ECM. Specifically, Legant et al. [18] made use of the inverse approach in conjunction with a linear elasticity theory to estimate tractions, while assuming that the strains in the matrix were small and the differences between the cell geometries in the stressed and relaxed states were negligible. However, cellular tractions can induce large deformations in the matrix with strain magnitudes more than 50% [18, 32]. Dong and Oberai [33] extended the inverse approach to account for finite deformations in the ECM, whose behavior was characterized by an exponential hyperelasticity model. This method was formulated in the reference (or unstressed) configuration of the ECM, and required the knowledge of the cell geometry in the unstressed state, which is often unknown due to cell lysis. In order to remedy this shortcoming, Song et al. [34] reformulated the problem in the current (or stressed) configuration of the ECM, thus only requiring the cell geometry in the stressed state. These authors also computed tractions exerted by 3T3 cells in dextran-based hydrogel matrices. More recently, Song et al. [35] generalized the inverse approach to estimate tractions in fibrous ECMs, making use of a Voigt-type (affine) homogenization model to explicitly account for the fibrous microstructure of the ECM.

The direct method has also been used to compute cellular tractions [15, 17, 36, 37, 32]. In this method, the measured discrete displacements are interpolated and extrapolated onto a continuous displacement field, which is differentiated to compute the strain field. The resulting strain field is then substituted into the constitutive equation to compute the stress and traction fields. Compared with the inverse approach, the direct method is computationally more efficient; however, its accuracy depends heavily on the quality of the strain field close to the cell surface. Given that spatial derivatives tend to amplify noise artifacts, this procedure requires a precise measurement of the displacement field.

In some studies cellular tractions have also been modeled as body forces that are distributed over the ECM volume [38], and are determined by solving an inverse problem. However, the approximation of surface tractions as body forces, especially if these forces are spread over a significant volume of the matrix, is questionable.

We are aware of only one computational method that is able to estimate cellular tractions while accounting for ECM heterogeneities [39]. In this approach, the measured dispalcements were differentiated to compute a strain field, which was then used in a discretized version of the equilbrium equations to determine the spatial distribution of the mechanical properties. Once these were determined, they were used to determine stresses and tractions by substituting the measured strain in the constitutive equation. While this approach is a notable extension to the standard 3DTM methods, it has a few drawbacks. First, it adopts an infinitesimal-strain, linear elasticity theory, which may be inadequate for most applications; further it is not clear how it might be extended to the finite-strain regime. Second, like most direct methods, it requires a precise measurement of the displacement field in order to arrive at an accurate estimate of strains and therefore stresses and tractions. In this manuscript, we develop a new approach for determining the modified ECM properties and tractions that address these issues. In addition, we provide a mathematical proof that it is possible to uniquely determine the mechanical properties and tractions from the data that is typically measured in the 3DTM experiment.

In this manuscript we employ a *compressible, hyperelastic* Neo-Hookean model [40] to describe ECM behavior, while allowing the material parameters (i.e., the two Lamé parameters) to be *nonuniform* in the ECM. We treat the material parameters as *unknowns* in regions proximal to the cell surface, acknowledging that enzymes synthesized by cells may have degraded the ECM thereby changing its mechanical properties [21]. In addition, we assume that matrix properties are unaffected by cell remodeling far from the cell surface, where the material parameters are uniformly distributed and can be determined from standard mechanical tests. Under these assumptions, and given a measured displacement field, we simultaneously determine the traction field at the cell-matrix interface, as well as the material parameters close to the cell surface.

The outline of the remainder of the manuscript is as follows. In Section 2 we present a mathematical formulation of the inverse problem of determining both tractions and material parameters. We show that the solution of this problem is unique, and present solvability conditions that are satisfied by perfect displacement data. In Section 3, we develop a novel optimization approach to solve the inverse problem. In Section 4, we evaluate the accuracy of our algorithm using synthetic data. In particular, we assess the sensitivity of our approach to noise, and quantify the error induced by neglecting the effect of ECM degradation. In Section 5 we summarize our findings and provide concluding remarks.

## 2. Problem formulation and uniqueness of the solution

In this section, we consider the problem of determining ECM degradation surrounding a cell, as well as the tractions exerted by the cell, given a single measured displacement field in the ECM in response to the tractions. We show that the solution to this problem is unique, provided that the given data satisfy certain solvability constraints.

### 2.1. Constitutive equations and ECM degradation

We consider a single cell embedded in a 3D ECM, which is taken to be isotropic and compressible [32, 41]. In particular, we employ a hyperelastic Neo-Hookean model [40] to characterize ECM behavior:

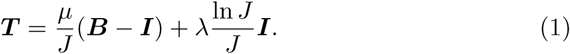

In the above equation, ***T*** is the Cauchy stress, ***B*** = ***FF*** ^*T*^ is the left Green-Lagrange stretch tensor, *J* = det(***F***) is the Jacobian, ***F*** is the deformation gradient tensor, and ***I*** is the second-order identity tensor. In addition, *μ* and *λ* are the two Lamé parameters of the ECM.

Since cells secrete enzymes (e.g., MMPs) that can locally degrade the ECM [21, 22], we assume that the two Lamé parameters are *nonuniform* across the degraded ECM. This effect can be represented by a degradation parameter *d*(***x***) (0 ≤ *d* < 1), so that

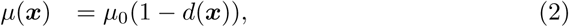

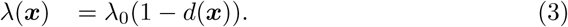

Here *μ*_0_ and *λ*_0_ are the Lamé parameters in the pristine state of the ECM (before it is degraded by cells), and ***x*** denotes a material point in the current configuration of the ECM. Note that *μ*_0_ and *λ*_0_ are known, and can be calibrated using standard mechanical tests (e.g., bulk rheology and indentation testings). On the other hand, *d*(***x***) is *unknown* a priori and needs to be determined (see below). In particular, ECM degradation depends sensitively on ECM composition, and is more severe for ECMs with high stiffness [23] and high proteolytic susceptibility [42]. Moreover, ECM degradation is confined within regions close to the cell surface (i.e., pericellular regions) [21], and diminishes progressively as we move away from the cell surface (*d* → 0). Note that we have used a single degradation parameter to describe the degradation of both Lamé parameters. The reasons for doing so will become evident below in Section 2.3 and the Appendix.

#### Remark

The hyperelastic Neo-Hookean model (1) accounts for the geometric nonlinearity due to large strains in the matrix; however, it does not include any form of material nonlinearities such as strain stiffening (or softening) at finite strains. For this reason, this model can be used to characterize the behavior of synthetic, bio-compatible hydrogels, such as the polyethylene glycol gels used in [18, 43]. However, it cannot be directly applied to realistic biological tissues, which often contain a significant proportion of fibers and exhibit highly nonlinear response at finite strains [44]. For those cases, more complicated models such as the fiber-based constitutive models (e.g., [35, 45, 46]) should be used.

### 2.2. Forward hyperelasticity problem

Next, we model the process of ECM relaxation after cell lysis. This process typically takes ∼ 45 mins [18, 34], and is commonly assumed to be quasi-static (e.g., [15, 17]). We further assume that ECM degradation mainly occurs during cell culturing (∼3 days [18]) before cell lysis, and that ECM properties do *not* vary significantly during the short period of ECM relaxation. Under these assumptions, it is sufficient to characterize ECM response using the hyperelasticity model in Section 2.1.

In particular, the balance of linear momentum for the ECM is given by

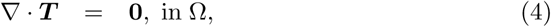

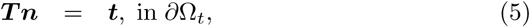

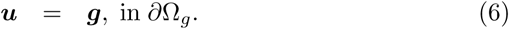

Here Ω is the spatial domain occupied by the ECM in the current configuration, *∂*Ω_*t*_ is the cell-ECM interface over which tractions are applied, and *∂*Ω_*g*_ is the ECM boundary over which displacement data ***u*** are prescribed. In our case this is the artificial boundary far away from the cell, where the displacement due to cellular tractions is assumed to have vanished. In addition, the operator **∇** denotes the spatial gradient (with respect to ***x***) in the current configuration, ***n*** is the unit outward normal on a given boundary, ***t*** is the traction field, and ***g*** is the prescribed Dirichlet data, which is set to zero in our problems.

Recall that the Cauchy stress ***T*** depends on the deformation gradient tensor ***F*** (see (1)), which in turn depends on the displacement field ***u*** via

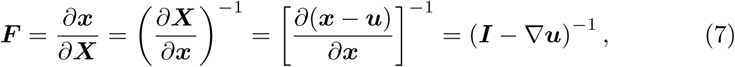

where ***X*** denotes a material point in the reference configuration. Given the degradation field *d*(***x***) (and hence the two Lamé parameters *μ*(***x***) and *λ*(***x***)) everywhere in the ECM, the displacement field ***u***(***x***) can be determined by solving equations (1)-(7). A similar method of solving finite-strain hyperelasticity problem in the current configuration can be found in [47].

*Weak form*. The forward problem (1)-(7) can be solved using the finite element method, which is based on the weak form, or the variational form of the problem.

More specifically, the weak form is stated as: find ***u*** ∈ 𝒮 such that

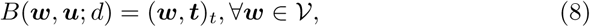

where

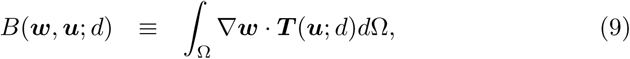

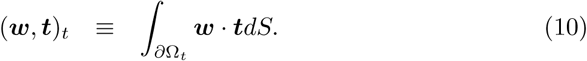

In the above equations, the function spaces 𝒮 and 𝒱 are defined as

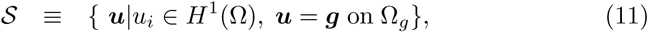

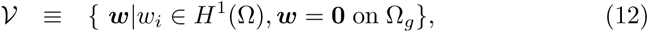

and *H*^1^ is the Sobolev space of square-integrable functions with square-integrable derivatives. Moreover, the dependence of ***T*** on the displacement field ***u*** and the degradation field *d* is made explicit in (9).

We solve the weak form (8) using the standard Galerkin finite element method [48], which approximates the infinite-dimensional functions space by their finite-dimensional counterparts. Moreover, we make use of the first-order Lagrangian basis functions associated with tetrahedral finite elements. This results in a set of nonlinear equations that can be solved by means of the Newton-Raphson method.

#### Remark

The forward hyperelasticity problem is formulated in the current (or stressed) configuration of the ECM, but not in the reference (or relaxed) configuration. This is motivated by the fact that the cell-ECM interface can be easily measured in the stressed state, while it is more challenging to do so in the relaxed state [18]. This is because the relaxed state is typically achieved by lysing the cell, which makes it difficult to locate the cell surface after cell lysis. It is possible to remove tractions while preserving cell integrity by using cell-contractility inhibitor (e.g., myosin inhibitor blebbistatin) [15]. In that case, the cell-ECM interface in the relaxed state can be measured, but the effectiveness of traction inhibition must be established to ensure complete stress relaxation. In any case, our formulation in the stressed configuration is particularly useful to model ECM relaxation in 3DTM.

### 2.3. Inverse hyperelasticity problem

In a 3DTM experiment, the displacement field in the ECM in response to cellular tractions is measured. Our objective is to recover the degradation parameter *d*(***x***) over the ECM volume, as well as the traction field ***t*** at the cell-ECM interface. Given that ECM degradation is expected to vanish (*d* = 0) at certain distances away from the cell [21], the inverse problem can be stated as follows: *Given the displacement field u*(***x***) *in* Ω, *and at least one point* ***x***_0_ *in* Ω, *where d*(***x***_0_) = 0, *find d*(***x***) *in* Ω *and* ***t***(***x***) *on ∂*Ω_*t*_ *such that equations (1)-(7) are satisfied*.

#### Theorem.

*If u*(***x***) *satisfies conditions (16) and (19) (see below), the solution to the above inverse problem is unique*.

*Proof*. Our proof follows the steps outlined in [49], where a similar result was outlined in the context of elastography, or elasticity imaging. Substituting the constitutive equations, (1)-(3), into the equation of equilibrium, (4), we have that

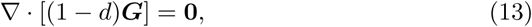

where the second-order tensor ***G*** is given by

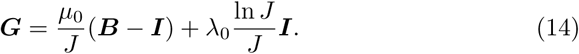

Note that given the measured displacement field ***u***, as well as the Lamé parameters (*μ*_0_ and *λ*_0_) for the pristine ECM, ***G*** is *known* everywhere in the ECM. Making use of the product role in equation (13), we arrive at

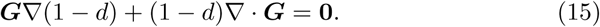

Assuming that 0 ≤ *d* < 1 and ***G*** is invertible, that is

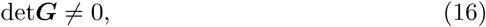

we can rewrite (15) as

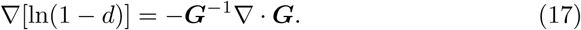

Integrating equation (17) from ***x***_0_, where *d* = 0, to any other point ***x*** ∈ Ω, we arrive at

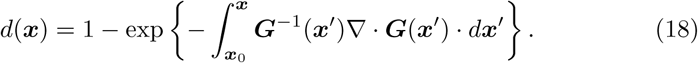

Having obtained *d*(***x***), we can determine the two Lamé parameters *μ*(***x***) and *λ*(***x***) everywhere in the ECM using (2) and (3), respectively. Finally, we can compute the Cauchy stress everywhere in the ECM via (1), and the traction ***t*** via (5).

Note that the integral in (18) is independent of the path between ***x***_0_ and ***x***, given that the integrand (i.e., the provided data) satisfies an integrability condition. In particular, for a simple connected region, such as a 3D ECM (since the ECM does not contain holes that penetrate all the way through it), this condition is

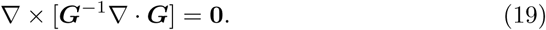

It is important to emphasize that performing the above operations requires two conditions, that is equations (16) and (19). In particular, (16) dictates that along a chosen integration path in (18), the Cauchy stress ***T*** at any point on that path must have *no* zero eigenvalues. As a result, in (16) an appropriate path must be selected to circumvent any singular points. Moreover, (19) places restrictions on the given data; if (19) is not satisfied, a single-valued solution of equation (17) for *d*(***x***) does *not* exist for that particular data. That is, the inverse problem is ill-posed.

#### Remarks

1. Equation (18) is an explicit expression for the degradation parameter in terms of the the tensor ***G***, which in turns depends on the displacement field. Thus, one may use this expression to evaluate *d* by approximating the integral in (18). However, this approach is likely to fail. This is because for real data which is corrupted by noise and by model discrepancy, the two conditions (16) and (19) may not hold, and the integral in (18) may diverge, or may yield path dependent values. Given this, in the following section we describe a regularized optimization-based approach that can handle noisy data that may be encountered in practice.
2. The degradation of the ECM has been assumed to be governed by a *single* degradation parameter *d*(***x***) (see equations (2) and (3)). This assumption implies that the Poisson’s ratio of the degraded ECM is identical to that of the pristine ECM, being unaffected by cell remodeling. However, it is possible that cells express enzymes (e.g., MMPs) that can alter the Poisson’s ratio of the ECM. In that case, it becomes necessary to reconstruct *λ*(***x***) and *μ*(***x***) independently. Unfortunately, the solution of this problem is generally *not* unique under the assumptions made in this section (see the Appendix for more details). Thus, such cases are not considered here and will be pursued in future work.
3. If there does not exist a point at which *d* = 0, or if the measured displacement field ***u*** has decayed to zero before reaching ***x***_0_ (recall that *d*(***x***_0_) = 0), it follows from (17) that the degradation field is determined up to an arbitrary multiplicative parameter (see similar results in [49]). Consequently, the stress and traction fields are also determined up to the same multiplicative parameter. However, ECM degradation is known to be concentrated near the cell surface [21] and, thus, the situation above is unlikely to occur.

## 3. Iterative optimization approach

In this section, we propose an iterative, optimization-based algorithm to simultaneously estimate the traction field at the cell-ECM interface and the degradation parameter in the ECM volume. For this purpose, we make use of a generalization of the inverse approach recently developed by [50]. In contrast to the method in Section 2.3, the iterative algorithm can handle noisy data that is encountered in practice.

### 3.1. Problem statement

As discussed in Section 2, we assume that ECM degradation occurs only at regions close to the cell surface, so that the ECM far from the cell is unaffected by cell remodeling and remains in its pristine state (*d* = 0, *λ* = *λ*_0_, *μ* = *μ*_0_). We further assume that the displacement field in the ECM is measured only at discrete bead locations, consistent with the standard experimental setup (e.g., [18, 34, 32, 20]).

We pose the inverse problem as a constrained optimization problem, where the goal is to find a traction field, ***t***, and a degradation field, *d*, that produce a displacement field that best matches the measured displacements. More specifically, the optimization problem is stated as follows: Given the displacement vectors measured at *K* bead locations, ***ũ***^*k*^ (*k* = 1, …, *K*), determine ***t*** and *d*, such that the objective function

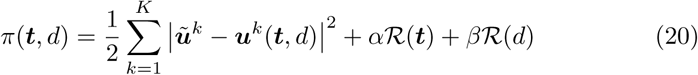

is minimized, under the constraint that the produced displacement field ***u*** satisfies the forward problem defined by (1)-(7) (or equivalently, the weak form (8)).

In equation (20), the first term is the data mismatch term, which evaluates the difference between the measured displacement vectors and the predicted displacement vectors (denoted by ***u***^*k*^) at the bead locations. Additionally, ℛ(***t***) and ℛ(*d*) are the regularization terms for ***t*** and *d*, respectively, and *α* and *β* are the associated regularization parameters. The regularization terms embed prior information about the unknown fields into the inverse problem. In most cases, these terms penalize unphysical spatial oscillations of the unknown fields, thus serving to ensure a certain smoothness of the solution to the ill-posed inverse problem. In this work, *H*^1^ semi-norm regularization [51] is used for both ***t*** and *d*:

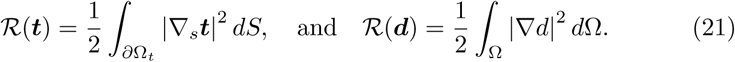

In the above expressions, ∇_*s*_ denotes the surface gradient operator such that ∇_*s*_***t*** = ∇ ***t***(***I*** − ***n*** ⊗ ***n***), keeping in mind that ***n*** is the unit outward normal to the cell-ECM interface.

In practice, the regularization parameters *α* and *β* are used to scale ℛ(***t***) and ℛ(*d*), respectively, so that the scaled regularization terms are balanced against the data mismatch term in the objective function. In this work, we make use of an extension of the L-curve method [52] to determine the optimal values of the regularization parameters. In a standard L-curve approach that handles a single regularization parameter, one needs to solve the inverse problem for different values of the regularization parameter, and plots the data mismatch term against the regularization term on a log-log scale, with the objective of identifying the data point that lies at the bend of this curve (i.e., the point of the maximum curvature). The regularization parameter that corresponds to this point is thought to optimally balance the data mismatch term and the regularization term in (20). The extension of this approach to multiple regularization parameters would require the generation of L-surfaces (see the results section), and the regularization parameters that correspond to the point lying at the bend (or the maximum curvature) of the surface are taken to be optimal.

#### Remarks

1. The formulation above does not compute tractions and material parameters directly from the measured displacement data. Instead, it aims to find the tractions and material parameters that would be most likely to produce the measured displacement field and satisfy some smoothness requirements associated with the regularization terms. As a consequence, this formulation relaxes the constraints of both (16) and (19), and can cope with noisy data.
2. Alternatively, one could determine the degradation parameter and the traction field sequentially. That is, one may first compute the degradation field by solving an inverse problem using an optimization approach, and then compute the stresses and tractions using the constitutive relation. However, the first step of this approach would require displacement data on the cell interface, which is often not available in a 3DTM experiment (e.g., [18, 34]). Therefore, we do not adopt this method. Instead, we determine the degradation parameter and the traction field simultaneously, requiring no displacement measurement on the cell interface.

### 3.2. Evaluation of the gradient

We solve the optimization problem using a quasi-Newton method, L-BFGS-B [53], which requires the gradient of the objective function *π* with respect to the traction field ***t*** and the degradation field *d*. The main difficulty lies in the fact that the displacement vectors ***u***^*k*^ (see (20)) depend on ***t*** and *d* implicitly through the constraint of the forward problem (8). Following the work of [50, 33, 34], we compute the gradient efficiently using an appropriate adjoint field, which can be derived by constructing a Lagrangian given by

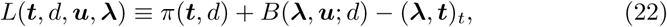

where ***λ*** ∈ **𝒱** plays the role of a Lagrangian multiplier field. The variation of *L* is given by

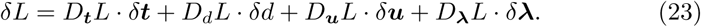

Here for a given function *F* that depends on the vector (or scalar) field ***v***, *D*_***v***_*F* · *δ****v*** denotes the Gateaux derivative defined by

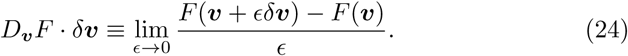

Setting *D*_***λ***_*L* · *δ****λ*** = 0, ∀*δ****λ*** ∈ 𝒱 in (23) leads to an equation for ***u***, i.e.,

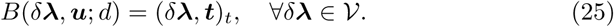

This equation implies that ***u*** satisfies the constraint of the forward problem (8). Under this condition, we have from (22) that

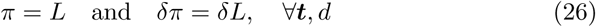

Further, we set *D*_***u***_*L* · *δ****u*** = 0, ∀*δ****u*** ∈ 𝒱 in (23) and arrive at

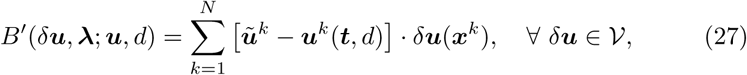

where the operator on the left hand side of (27) is given by

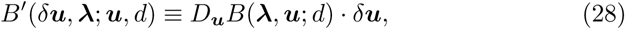

and ***x***^*k*^ denotes the location of the *k*th bead. We note that (27) leads to a well-posed linear problem for the Lagrangian multiplier field ***λ***. In particular, the left hand side of this problem is the adjoint of the linearization of the original forward problem (8). Therefore, we refer to (27) as the adjoint problem, and ***λ*** the adjoint field. We also note from (27) that the solution of ***λ*** is driven by the data mismatch term.

With ***u*** and ***λ*** given by the solution of (25) and (27), respectively, we have from (23) and (26) that

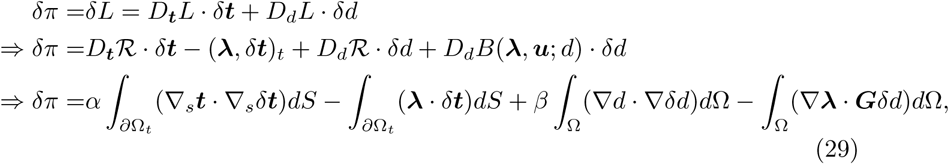

where we recall that the second-order tensor ***G*** is defined by (14), and *α* and *β* are the regularization parameters (see (20)) associated with ***t*** and *d*, respectively. Similar strategies have been used to solve the inverse problems for the spatial distribution of material parameters (e.g., [54, 55, 56]), although in a different context.

In summary, given the measured displacements ***ũ***^*k*^ (*k* = 1, …, *K*), as well as the mechanical properties of the pristine ECM (*λ*_0_ and *μ*_0_), we can reconstruct the traction field, ***t***, and the degradation field, *d*, as follows:

1. For given guesses of ***t*** and *d*, solve equation (25) to determine the primal field ***u***.
2. Solve equation (27) to determine the adjoint field ***λ***.
3. Employ the primal and adjoint fields to determine the objective function and the gradient vector using (20) and (29), respectively.
4. Utilize the L-BFGS-B algorithm to obtain updated guesses for ***t*** and *d*. Then, steps 1-4 are repeated until convergence. The L-BFGS-B iterations are considered to be converged when the relative change in the data mismatch term over the last five iterations is less than 1 × 10^−8^.

In practice, we work with finite-dimensional counterparts of these fields. As already mentioned, we make use of first-order Lagrangian basis functions and tetrahedral finite elements. Note that the traction field and its variations are defined only on the cell-ECM interface, while all the other fields are defined in the ECM volume. Also note that ***t*** = **0** and *d* = 0 are used as the initial guesses.

In summary, an iterative optimization algorithm has been developed to quantify in 3D both the cellular tractions and ECM degradation. Thus, the new algorithm will be heretofore referred to as the 3D Traction and Degradation Microscopy (3DTDM) algorithm. Remarkably, the new algorithm requires the same input as a standard TM algorithm, which includes the cell geometry in the stressed state, a single measured displacement field in the ECM, as well as the mechanical properties of the pristine ECM. For this reason, it can be directly applied to experimental data acquired in a standard TM experiment. In addition, the degradation field predicted by 3DTDM represents the biochemical imprint of cells on the ECM, and is expected to contain useful information about the biochemical activities of cells.

## 4. Results

In this section, we validate the new 3DTDM algorithm *in silico* by quantifying its error in recovering “known” degradation and traction fields. We also examine the effect of neglecting ECM degradation on traction measurements.

### 4.1. Problem setup

#### 4.1.1. Generating synthetic data

We assume that cell geometry is known in the stressed state (prior to cell lysis), consistent with standard 3DTM measurements. Given this, a chosen degradation field 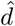 in the ECM, and a chosen traction field 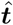 at the cell-ECM interface, we compute the true displacement field ***û*** sampled at the bead locations by solving a forward problem (equation (8)). Then, we add noise to these displacement vectors to mimic realistic measured displacements in the experiment. In particular, the measured displacement vectors ***ũ***^(*k*)^ (*k* = 1, …, *K*) are given by

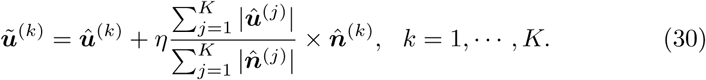

Here 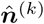 is a random vector, with its three components given by 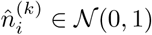, *i* = 1, 2, 3, and 𝒩(0, 1) is the standard normal distribution. Moreover, *η* represents the level of noise in the synthetic measured displacement field. In this work, we choose *η* = 1% and 5%, which are thought to be realistic in a TM experiment [57].

#### 4.1.2. Inverse problem and error analysis

Making use of the synthetic displacements (30) as input, we determine the degradation and traction fields using the 3DTDM algorithm developed in Section 3. Moreover, in order to assess the error induced by neglecting ECM degradation, we use the same synthetic displacements (30) as input, but determine only the traction field ***t***, while incorrectly assuming that no degradation occurs (*d* = 0). The latter approach is equivalent to a standard 3DTM approach.

Solving the inverse problem yields a reconstructed degradation field, *d*, and a reconstructed traction field, ***t***. Making use of these fields, together with the true degradation field 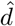 and the true traction field 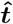, we assess the performance of the inverse algorithms by quantifying the relative error *e*(*d*) and *e*(***t***):

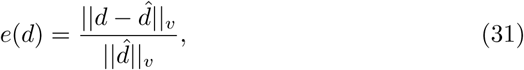

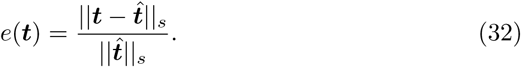

Here ‖ · ‖_*v*_ denotes the *L*_2_ norm of a given field defined in the ECM volume Ω, while ‖ · ‖_*s*_ denotes the *L*_2_ norm of a given field defined on the cell-ECM interface *∂*Ω_*t*_. The metrics defined by equations (31) and (32) contain useful information. For 3DTDM, they represent the reconstruction error induced by noise in the measured displacement; for 3DTM, they also include the error induced by neglecting ECM degradation. Note that by definition, *e*(*d*) = 100% for 3DTM.

We also examine the capability of a given algorithm to recover the measured displacement field. For this purpose, we evaluate the difference between the measured displacements ***ũ***^*k*^ and the predicted displacements ***u***^*k*^ (*k* = 1, …,*K*). In particular, we define the relative error *e*(***u***) as follows:

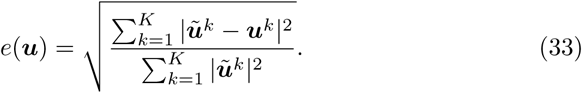

We remark that *e*(***u***) in (33) is especially useful for assessing the accuracy of the inverse algorithm in real experiments, in which case (31) and (32) cannot be used, since the true degradation field 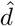 and the true traction field 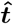 are *not* known.

### 4.2. Numerical examples

#### 4.2.1. Generating synthetic data

Here we consider a geometrically accurate, neuronal cell model (Fig. 1). The geometric model and finite element mesh (Fig. 1a) are generated from a set of Z-stack microscopy images available at [58]. The mesh contains 10,673 vertices and 54,587 tetrahedral elements. The cell is approximately 10 × 10 × 10*μ*m, embedded within a cube with sides of length 80*μ*m. Since the outer boundaries of the domain are far from the cell surface, cell-generated displacements are assumed to have vanished on those boundaries.

**Figure 1:**
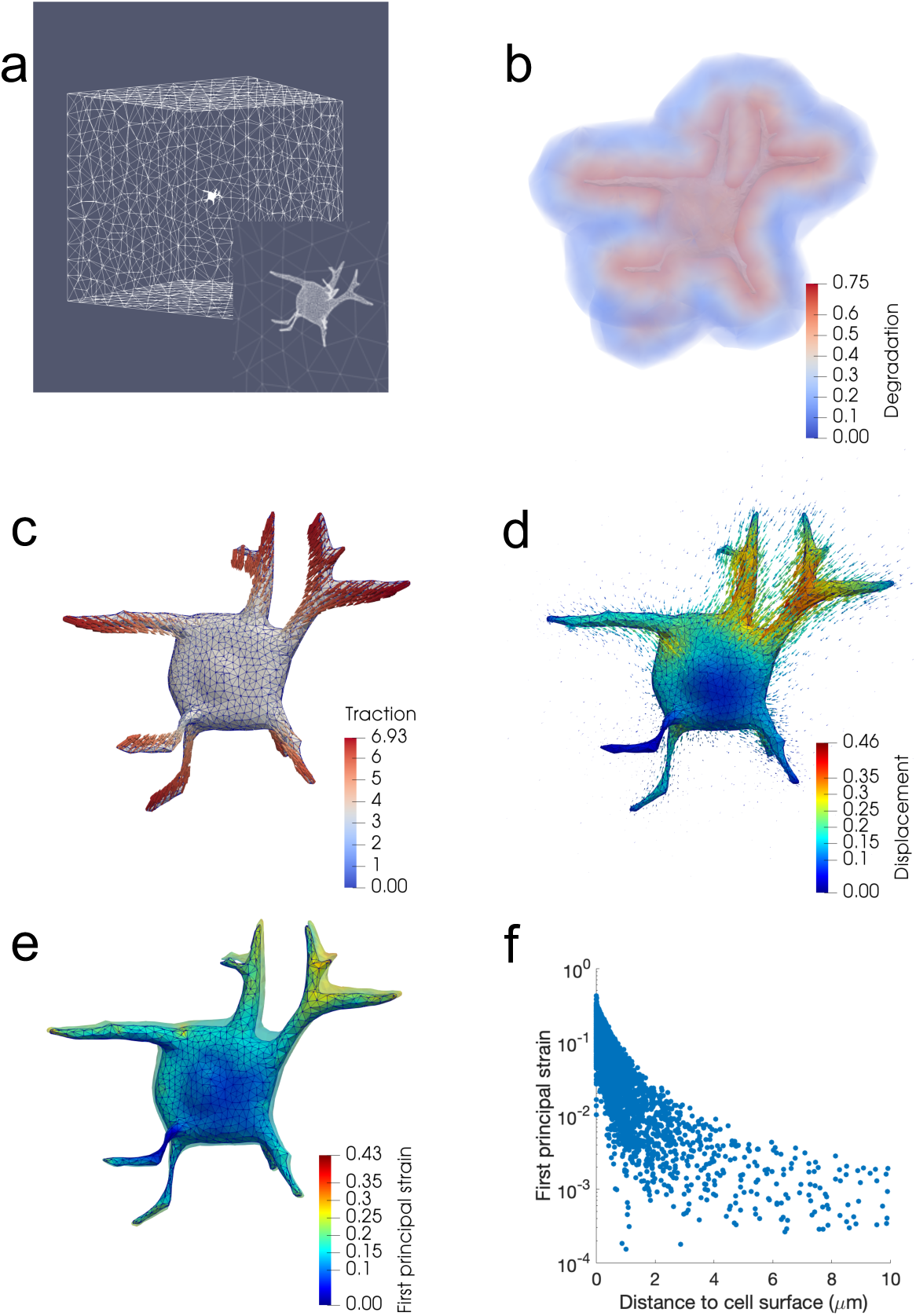
Forward problem results for a neuronal cell. (a) Finite element mesh, with the inset showing mesh refinement on the cell surface. (b) 3D rendering of the simulated degradation field within 2*μ*m of the cell surface. (c) Simulated traction field (kPa) applied to the cell. (d) Displacement field (*μ*m) in response to applied tractions. The magnitude of the displacement field on the cell surface is also color-coded. (e) First principal Eulerian strain on the cell surface. The surface of the cell in the unstressed configuration is also displayed as a semi-transparent outline around the stressed configuration. (f) First principal Eulerian strain at mesh vertices as a function of distance from the cell surface.

As discussed above, ECM degradation is concentrated within domains near the cell, and is more severe for ECMs with high stiffness and high proteolytic susceptibility. Motivated by this, we assume that degradation occurs only in regions within 2*μ*m of the cell surface (Fig. 1b), being most severe at the cell-ECM interface and diminishing progressively as we move away from the cell surface. To demonstrate the effect of ECM degradation, we confine our attention to cases where the ECM is significantly degraded. In particular, we set *d* = 0.75 at the cell-ECM interface, and *d* = 0 for regions beyond 2*μ*m of the cell surface. For domains within 2*μ*m of the cell surface, the values of *d* are linearly interpolated between 0.75 and 0, as a function of the distance from the cell surface. More specifically, the degradation field *d*(***x***) is defined as:

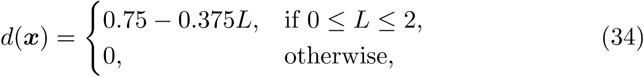

where *L* denotes the distance between a point ***x*** and the cell surface. With this choice, the pristine ECM is four times stiffer than the degraded ECM at the cell surface, representing a typical case with significant degradation [59]. Unfortunately, we are not aware of any experimental results showing the detailed 3D distribution of the degradation field, and *d*(***x***) in (34) may not be realistic. However, for the purpose of demonstrating the capability of our algorithm *in-silico*, we are satisfied with the simple choice in (34). Further, we set the Lamé parameters of the pristine ECM to be *μ*_0_ = 20kPa and *λ*_0_ = 80kPa, yielding a Poisson’s ratio of 0.4. These material parameters are thought to be appropriate for ECMs with high stiffness, and are within the range of the typical values used for TM studies (e.g., [57, 41, 60]).

Motivated by realistic tractions measured in previous studies [18, 34], the traction field is applied such that its magnitude is largest at the tips of the protrusions, and decreases gradually towards the cell body (Fig. 1c). Moreover, the direction of the traction is roughly pointed to the cell center (Fig. 1c), emulating the behavior of a contractile cell.

Given a chosen degradation field (Fig. 1b), and a chosen traction field (Fig. 1c), we solve a forward problem to determine the resulting displacement field (Fig. 1d). These displacements lead to significant local strains in the ECM (Fig. 1e and f), where the first principal value of the Eulerian strain, ***e*** = 1/2(***I*** − ***B***^−1^), can be as large as 43%. Additionally, we reconstruct the cell surface in the unstressed state by warping the stressed cell surface by the displacement vector (i.e., ***X*** = ***x*** − ***u***). The result is shown as a transparent surface in Fig. 1e. As can be seen, the cell geometry in the unstressed and stressed states are noticeably different, particularly at the top two protrusions, suggesting that nonlinear modeling should be used to estimate degradations and tractions.

The bead locations are taken to be all the mesh vertices (excluding those on the cell surface) within a sphere of radius 25*μ*m centerd at the centroid of the domain. This choice yields a total of 6,971 beads, which is comparable to the bead number in a standard 3DTM experiment [18, 34]. In addition, the above choice is consistent with the experimental protocol where most of the beads that are near the cell surface are tracked. Further, we add noise to displacement vectors at the bead locations (equation (30)) to create the “measured” displacement field, which will be used as input for the inverse problem.

#### 4.2.2. Inverse problem and error analysis

For 3DTDM, the unknown variables are taken to be the nodal values of the traction vector ***t***, as well as of the degradation parameter *d* within 2*μ*m of the cell surface. For regions beyond 2*μ*m of the cell surface, *d* is set to 0 and considered to be *known*, since no degradation occurs in those regions.

##### Simultaneous reconstructions

We first examine the accuracy of 3DTDM in recovering both the degradation and traction fields. To solve the inverse problem, we need to determine the optimal values of the regularization parameters, that is *α* and *β* in equation (20). For this purpose, we make use of the L-surface approach, as mentioned in Section 3.1. In particular, we generate the L-surfaces by solving the inverse problems for a range of *α* and *β*, and plot the data mismatch term (the first term in (20)) as a function of the regularization terms ℛ(***t***) and ℛ(*d*). The resulting L-surfaces for 1% and 5% noise are shown in Fig. 2. We attempt to identify the points that are close to the bend (or the maximum curvature) of the L-surfaces (see the black dots in Figs. 2a and b) [50]. The regularization parameters that correspond to those points are taken to be optimal, generalizing the idea of a standard L-curve approach [52]. In particular, the optimal regularization parameters are: *α* = 1 × 10^−5^, *β* = 5 × 10^−5^ for 1% noise, and *α* = 1 × 10^−4^, *β* = 2.5 × 10^−4^ for 5% noise. Note that it is possible to improve our selection procedure by generating more points on the surface (i.e. running more simulations for different regularization parameters) and pursuing a quantitative means of assessing the curvature at each simulation. However, the L-surface approach is heuristic so the extra effort involved is difficult to justify.

**Figure 2:**
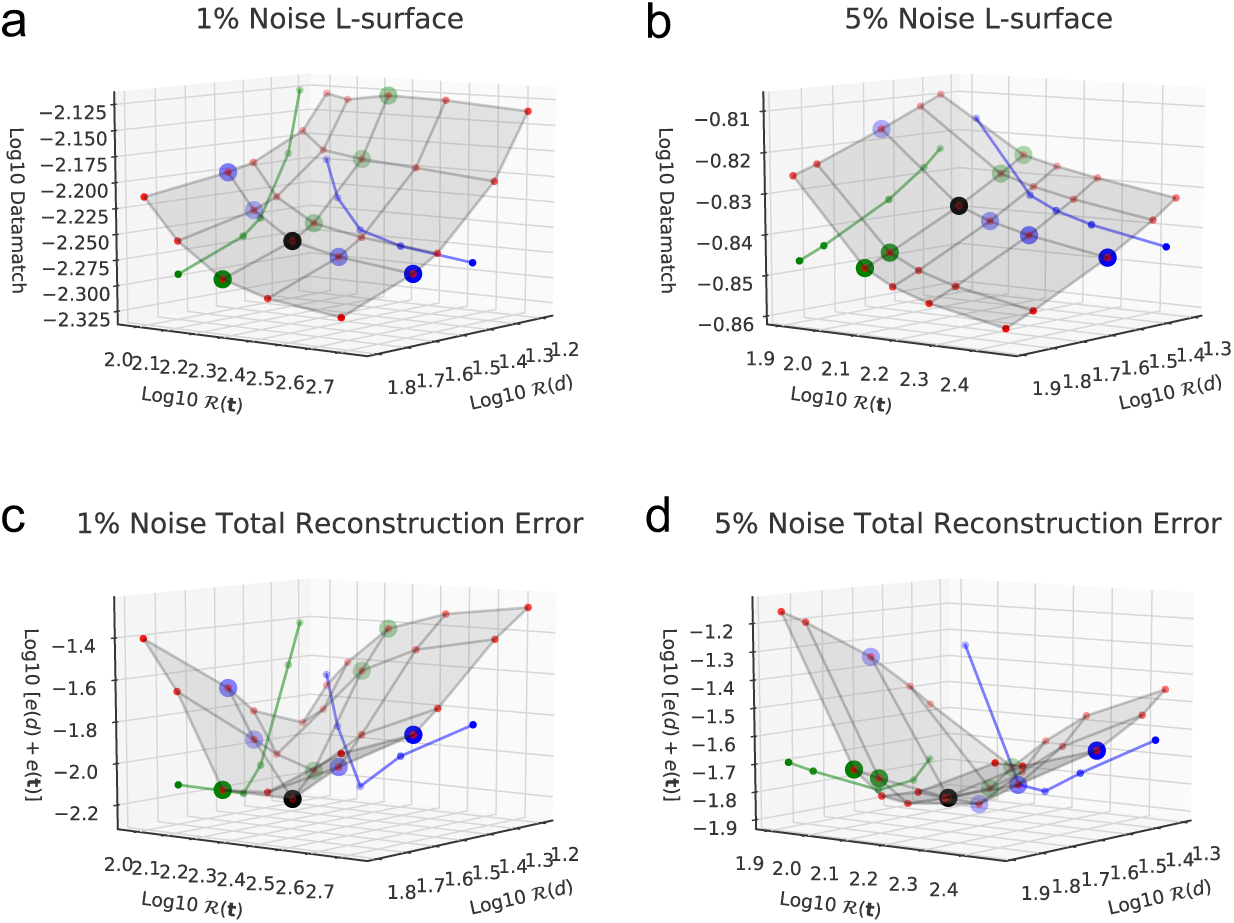
L-surfaces plots obtained by solving the inverse problem for a range of regularization parameters (*α* and *β* defined in (20)) for (a) 1% noise, and (b) 5% noise. The simulations that are considered to be at the “bend” of the surfaces are marked by the black dots, which correspond to (a) *α* = 1 × 10^−5^, *β* = 5 × 10^−5^, and (b) *α* = 1 × 10^−4^, *β* = 2.5 × 10^−4^. Projections of the L-surfaces that correspond to fixed values of the regularization parameters are displayed on the “walls” of the graphs. In particular, the green and blue curves correspond, respectively, to (a) *α* = 1 × 10^−5^, and *β* = 5 × 10^−5^; (b) *α* = 1 × 10^−4^, and *β* = 2.5 × 10^−4^. The total reconstruction error, *e*(***t***) + *e*(*d*), for each point displayed on the L-surfaces is used to construct the error surfaces for (c) 1% noise, and (d) 5% noise.

To illustrate the effectiveness of the L-surface approach, we display in Figs. 2c and d the total reconstruction error, *e*(***t***) + *e*(*d*), that corresponds to each point on the L-surfaces in Figs. 2a and b. Note that in practice, the total reconstruction error is *unknown*. Therefore, in the spirit of fairness, we selected the points in the “bend” of the L-surfaces, as marked by black dots in Figs. 2a and b, prior to creating the error plots. We can see from Figs. 2c and d that for both noise levels, the total reconstruction error that corresponds to the black dot is near the true minimal value, suggesting that the L-surface method is useful.

Figure 3 displays 3DTDM results for the degradation field *d*, traction field ***t***, and the local traction error 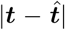 for both 1% and 5% noise. The exact degradation and traction fields are also shown for comparison (Figs. 3a, d, and g). We observe from Figs. 3a-f that for both noise levels, 3DTDM recovers the degradation field reasonably well. In particular, the degradation reconstruction is most sensitive to noise in regions near the cell surface, where increasing the noise from 1% to 5% leads to greater error in those regions (e.g., compare Fig. 3e with Fig. 3f). In addition, 3DTDM recovers the traction field fairly accurately (Figs. 3g-i), with traction errors being lower for 1% noise than for 5% noise (Figs. 3j and k). Note that for the 5% noise case, the traction errors are noticeable at the cell body (Fig. 3k), although tractions are rather small at those regions (Fig. 3g). This could be due to the fact that the “measured” displacements near the cell body are dominated by noise, since the original signal is weak.

**Figure 3:**
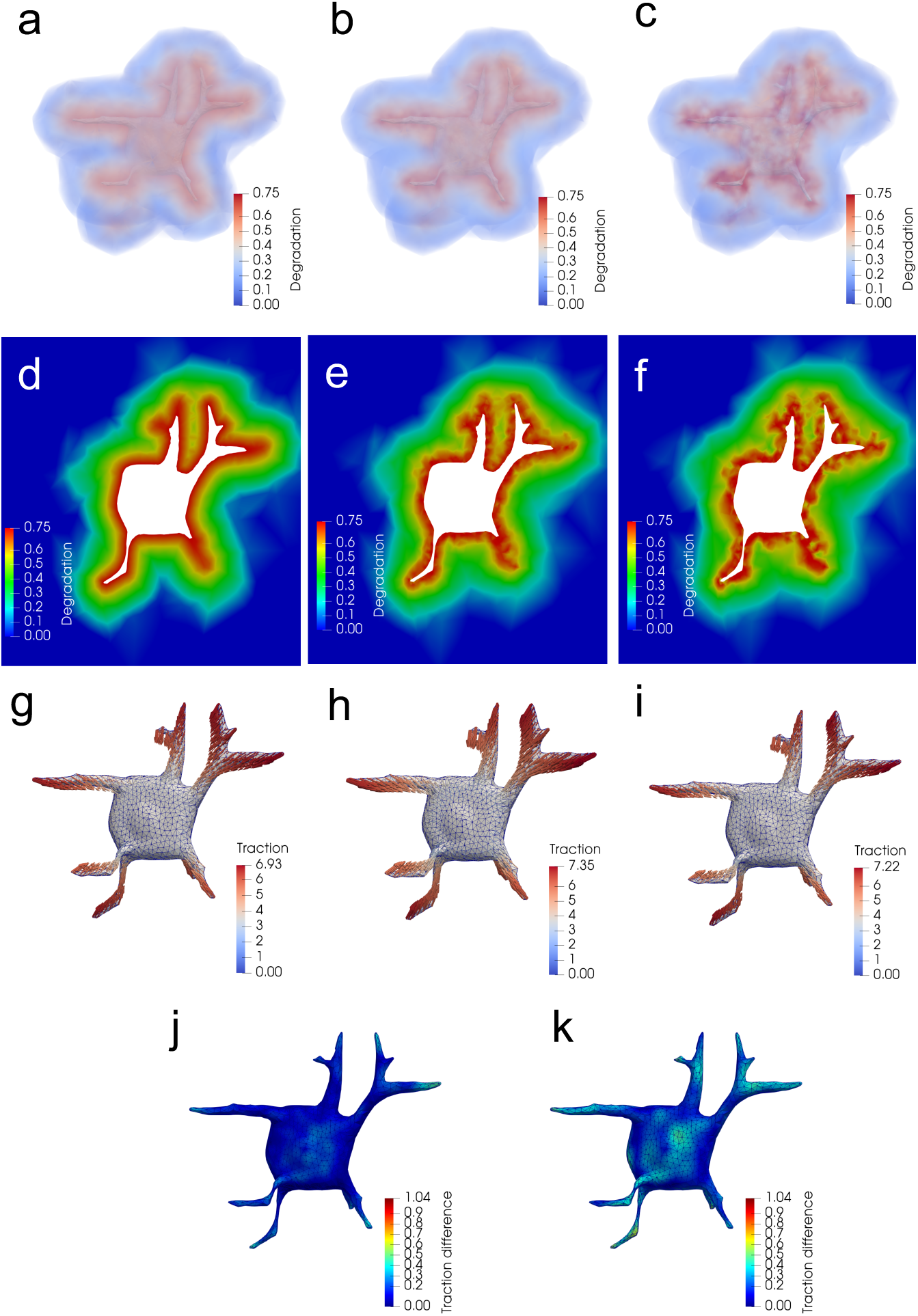
Inverse problem results obtained by simultaneously reconstructing the degradation field, *d*, and the traction field, ***t***, for a neuronal cell with 1% and 5% noise. Results are shown for the 3D rendering of the degradation field *d*: (a) exaction solution, (b) 1% noise, (c) 5% noise; Degradation field *d* on a 2D cross section of the ECM: (d) exact solution, (e) 1% noise, (f) 5% noise; Traction field (kPa): (g) exact solution, (h) 1% noise, (i) 5% noise; Local traction error 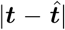 (kPa): (j) 1% noise, (k) 5% noise.

##### Traction-only reconstructions

Next, we investigate the error induced by neglecting ECM degradation. To this end, we examine the accuracy of a standard 3DTM algorithm in estimating cellular tractions. To prepare for the inverse problem, we first determine the optimal regularization parameter *α* for the traction field using the standard L-curve approach [52]. The optimal regularization parameters are taken to be those associated with the points that are close to the maximum curvature of the L-curves (see the black points in Fig. 4). Note that the L-curves for both noise levels appear to be rather similar. This is because for traction-only reconstructions, data mismatch is primarily induced by neglecting ECM degradation (see below), and not so much by measurement noise. In particular, *α* = 5 × 10^−4^ is used for both 1% and 5% noise.

**Figure 4:**
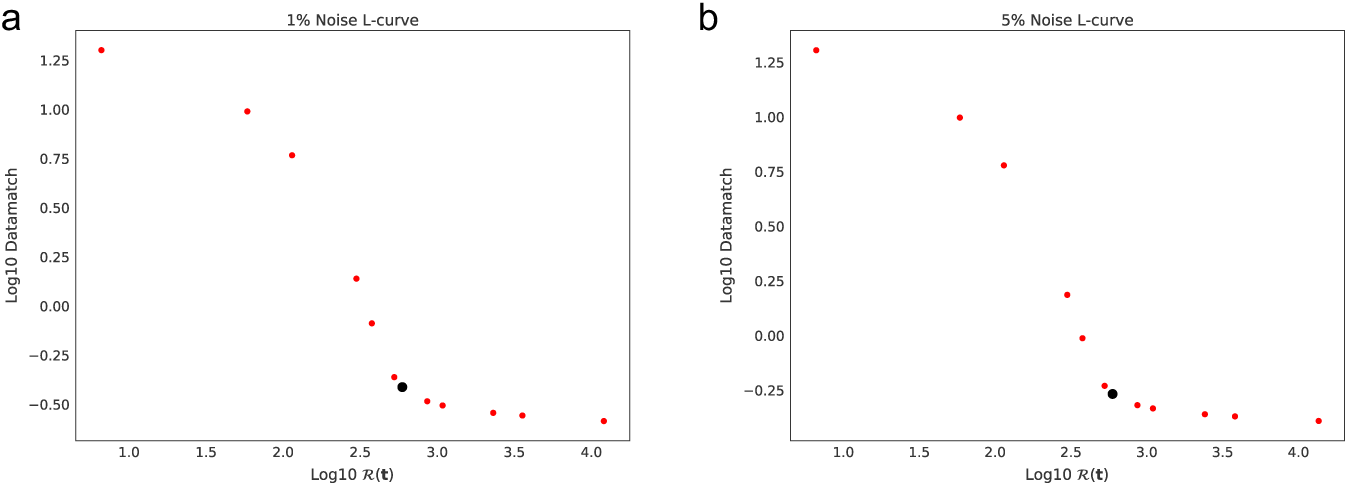
L-curves plots obtained by solving the inverse problem for a range of regularization parameters *α* defined in (20)) for (a) 1% noise, and (b) 5% noise. The simulations that are considered to be at the “bend” of the curves are marked by the black dots, which correspond to *α* = 5 × 10^−5^ for both noise levels.

Figure 5 shows 3DTM results for the traction field ***t*** (Fig. 5a), and the first principal Eulerian strain (Fig. 5c) with 1% noise. (The 3DTM results for 5% noise are similar to those for 1% noise and are not shown). For comparison, the corresponding exact fields are also displayed in Figs. 5b and d. We can see from Fig. 5a and b that 3DTM drastically overestimates the traction field, and the maximum predicted traction is almost two times the exact value. At the same time, 3DTM significantly underestimates the strain on the cell surface, where the recovered maximum strain is roughly 40% lower than the exact value (compare Figs. 5c and d).

**Figure 5:**
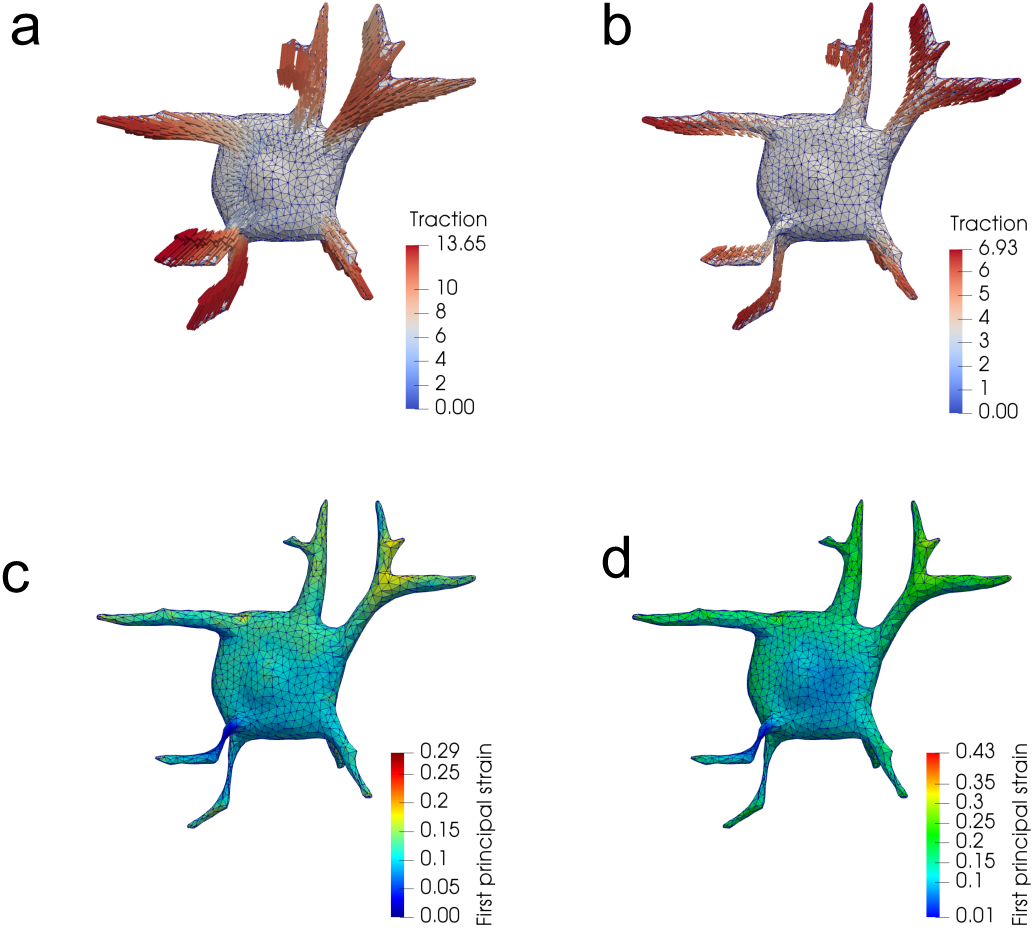
Inverse problem results obtained by reconstructing only the traction field, ***t***, for a neuronal cell with 1% noise. The corresponding exact solutions are also shown for comparison. (a) Reconstructed traction field (kPa), (b) Exact traction field (kPa), (c) Reconstructed first principal Eulerian strain, and (d) Exact first principal Eulerian strain.

The above results can be explained by examining Fig. 6, which shows the true displacement field (Fig. 6a), as well as the displacement field recovered by 3DTM (Fig. 6b), on a 2D cross section of the ECM. Specifically, the true displacement field decays rapidly as we move away from the cell surface (Fig. 6a). This is because the ECM is severely degraded near the cell, and is weaker in transmitting forces. However, this phenomenon cannot be captured by 3DTM, which neglects ECM degradation thus predicting a slower displacement decay (Fig. 6b). When solving an inverse problem, 3DTM attempts to match the true decay by underestimating the displacement (and strain) near the cell and overestimating them away from the cell (Fig. 6b). 3DTM also uses an ECM that is much stiffer than the true ECM at the cell-ECM interface (keeping in mind that 3DTM incorporates no ECM degradation), thus predicting overall larger tractions, which are roughly the product of stiffness with strain.

**Figure 6:**
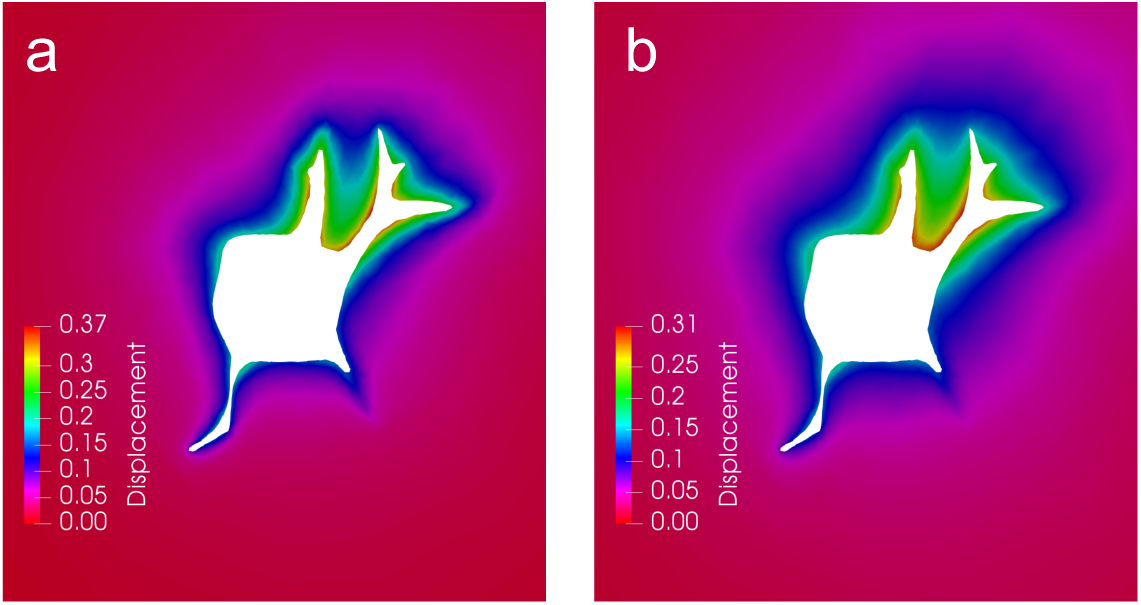
Magnitude of the displacement field (*μ*m) on a 2D cross section of the ECM. (a) Exact results, and (b) Reconstructed results for 1% noise, where ECM degradation is neglected.

Note that for the cases considered above, where the ECM degradation is significant, a direct TM method is likely to introduce even larger errors. This is because it directly uses the measured displacement field to compute strains, which are substituted into the constitutive relation for a *pristine* ECM to evaluate stresses and tractions. As a consequence, the error in tractions would be in direct proportion to the degree of ECM degradation at the cell-ECM interface.

###### Remark

Our results suggest that cell-induced ECM degradation could give rise to rapid displacement decay and thus short-range force transmission (Fig. 6a). On the other hand, previous studies have shown that for fibrous ECM, cell-induced fiber realignment leads to ECM stiffening, as well as to slow displacement decay and long-range force transmission (e.g., [35, 61, 62]). Taken together, these results suggest that cells can modulate the range of force transmission in fibrous ECMs through two competing mechanisms—ECM degradation and fiber realignment. Thus, it is of great interest to extend the current method to measure tractions in fibrous ECMs, accounting for both fiber degradation and fiber realignment.

##### Error analysis

Table 1 summarizes the reconstruction errors for both 3DTDM and 3DTM with 1% and 5% noise. It is observed that for both noise levels, 3DTDM recovers the degradation and traction fields accurately. In particular, for 5% noise it yields 6.26% error in tractions and 21.7% error in degradation. Additionally, 3DTDM matches the measured displacements very well, yielding a data mismatch of 4.59% for 5% noise. In contrast, 3DTM yields about 103% traction errors for both noise levels. As discussed above, these errors are induced by the failure of incorporating ECM degradation. These results suggest that it is important to account for ECM degradation in order to accurately recover cellular tractions, particularly for cases with significant ECM degradation (e.g., ECMs with high proteolytic susceptibility and high stiffness). Thus, the new 3DTDM approach can extend the range of context where traction microscopy can be appropriately applied.

**Table 1:** Reconstruction errors for the degradation field *d*, traction field ***t***, and displacement field ***u***, as defined by equations (31), (32), and (33), respectively. Results are shown for both 3DTDM and 3DTM at two noise levels *η* = 1% and 5%.

|  | 3DTDM |  | 3DTM |  |
| --- | --- | --- | --- | --- |
| Error | $\eta = 1\%$ | $\eta = 5\%$ | $\eta = 1\%$ | $\eta = 5\%$ |
| $e(d)$ (%) | 15.4 | 21.7 | 100 | 100 |
| $e(\mathbf{t})$ (%) | 3.68 | 6.26 | 102.9 | 103.0 |
| $e(\mathbf{u})$ (%) | 0.881 | 4.59 | 7.56 | 8.91 |

## 5. Concluding Remarks

In this work a novel computational approach was developed to determine tractions exerted by cells fully encapsulated in 3D extracellular matrices (ECM), while accounting for cell-induced ECM degradation. The behavior of the ECM was characterized by a Neo-Hookean hyperelasticity model, with Lamé parameters corrupted by a single parameter that quantified ECM degradation. The degradation was assumed to be confined within regions near the cell surface. It was most severe at the cell-ECM interface and diminished progressively with increasing distance from the cell.

The problem was posed as an inverse problem where the goal was to determine the degradation field and the traction field that were consistent with a measured displacement field. First, the feasibility of quantifying both the traction and degradation fields was demonstrated by demonstrating the existence and uniqueness of the solutions, under certain solvability conditions. Then, an iterative algorithm, which was referred to as the 3D Traction and Degradation Microscopy (3DTDM), was developed to simultaneously determine the degradation and traction fields. The inverse problem was formulated as an optimization problem, with the objective of minimizing the difference between a measured and predicted displacement field, under the constraint that the predicted displacement field satisfied the equations of equilibrium. The problem was solved by means of a gradient-based minimization approach, and the gradient was evaluated efficiently using an adjoint method.

The 3DTDM algorithm was validated *in silico* by quantifying its error in recovering known, synthetic degradation and traction fields for a neuronal cell. For a measured displacement field with realistic noise (e.g., 5%), the algorithm was able to recover both the degradation and traction fields fairly accurately. Furthermore, it was found that neglecting ECM degradation tended to over-estimate cellular tractions, highlighting the importance of incorporating ECM degradation to accurately measure these tractions.

The 3DTDM approach developed in this work can be improved in several ways. For instance, the degradation of the bulk and shear modulus of the ECM could be considered independently to more accurately capture ECM degradation. However, as shown in the Appendix, the solution of the inverse problem is non-unique. Further investigations are warranted to address this challenging issue. In addition, while the hyperelastic Neo-Hookean model can be used for synthetic, bio-compatible hydrogels, it cannot be used for realistic fibrous ECMs, which often exhibit highly nonlinear response at finite strains. To capture the complex behavior of those materials, more advanced constitutive models, such as the fiber-based homogenization models (e.g., [35, 45]), should be used. We believe that the ability to recover tractions in 3D fibrous ECMs, while accounting for both fiber degradation and fiber realignment, will be a significant step forward.

To the best of the our knowledge, the 3DTDM approach described in this work is the first method that is able to measure both cellular tractions and ECM degradation in a finite-strain setting. Remarkably, it requires the same input data as needed in a standard 3D traction microscopy, making it applicable to a broad range of experimental settings. In addition to accurately measuring cellular tractions, the 3DTDM approach can also provide useful information about the biochemical imprint of cells on the ECM. Thus, it may serve as a robust tool to study both the mechanical and biochemical activities of cells in important cellular processes such as migration and proliferation.

## Acknowledgments

This material is based upon work supported in part by the National Institute of Health under Grant Numer 1R41CA224898-01. The authors declare no conflict of interest.

Sandia National Laboratories is a multimission laboratory managed and operated by National Technology and Engineering Solutions of Sandia, LLC, a wholly owned subsidiary of Honeywell International Inc., for the U.S. Department of Energy’s National Nuclear Security Administration under contract DE-NA0003525.

This paper describes objective technical results and analysis. Any subjective views or opinions that might be expressed in the paper do not necessarily represent the views of the U.S. Department of Energy or the United States Government.

## Appendix

In this Appendix, we consider the inverse problem of determining the spatial distribution of the two Lamé parameters (*λ* and *μ* in (1)) for a compressible, Neo-Hookean ECM, given a single measurement of the displacement field in the ECM. In particular, we demonstrate using a specific example that the solution of this problem is *not* unique. This approach build upon similar results presented in papers dedicated to elastography, or elasticity imaging [63, 64].

### A1. Inverse problem

For simplicity, and without loss of generality, we consider the case with infinitesimal strains (i.e., |**∇*u***| ≪ 1). Neglecting the higher-order terms, we can easily show that equation (1) reduces to the linear Hooke’s law:

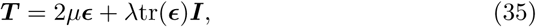

where ***ϵ*** denotes the infinitesimal strain tensor given by

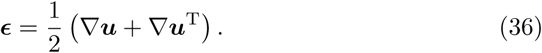

In addition, we assume that ECM degradation is confined within a region proximal to the cell surface, so that the ECM is in its pristine state (i.e., *λ* = *λ*_0_ and *μ* = *μ*_0_) far from the cell surface.

Thus, we consider the following inverse problem: *given the displacement field* ***u***(***x***) *in the ECM occupying a domain* Ω, *and at least one point* ***x***_0_ *in the ECM where λ* = *λ*_0_ *and μ* = *μ*_0_, *find μ*(***x***) *and λ*(***x***) *in* Ω *such that equations (4), (35), and (36) are satisfied*.

#### A2. General solution

To solve the inverse problem, we substitute (35) into (4) and arrive at

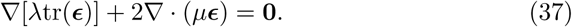

Note that given ***u***, we can compute ***ϵ*** using (36). Thus we may consider ***ϵ*** known. We eliminate *λ* by taking the curl of (37) to arrive at

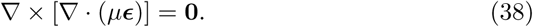

Once *μ* is solved from (38), we can determine *λ* straightforwardly using equation (37) [49]. More specifically, we integrate (37) from ***x***_0_, where *λ* = *λ*_0_, to any other point ***x*** ∈ Ω, and arrive at

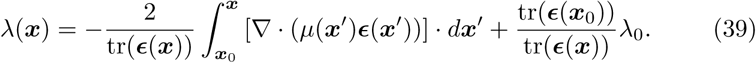

Note that in (39) we have assumed that tr(***ϵ***(***x***)) = **∇** · ***u***(***x***) ≠ 0, which requires that the deformation at ***x*** must induce local changes in the ECM volume.

Now we go back to equation (38), which is an overdetermined system having three equations for only one unknown. In general, the solution of an overdetermined set of partial differential equations may not exist, but it may also permit multiple solutions [64]. Since ***ϵ*** in (38) represents the response of an actual ECM, which possesses two Lamé parameters *μ* and *λ*, we are guaranteed that at least one solution exists. We demonstrate below using a simple example that the solution of (38) is generally *not* unique.

### A3. Special case of spherical symmetry

Here we consider a special case where the provided displacement field ***u***(***x***) exhibits spherical symmetry. In particular, ***u***(***x***) and the associated ***ϵ***(***x***) can be written as

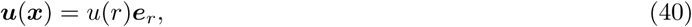

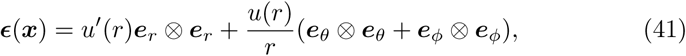

where ***e***_*r*_, ***e***_*θ*_, and ***e***_*ϕ*_ (0 ≤ *r*, 0 ≤ *θ ≤ π*, 0 ≤ *ϕ* ≤ 2*π*) are the three basis vectors in the spherical coordinate system.

Then, we can easily verify that any choice of

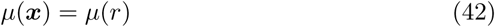

is a solution of equation (38). To see this, for any *μ* given by (42), we have that

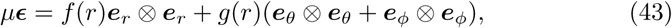

where *f* = *μu*′ and *g* = *μu/r* are functions of only *r*. Taking the divergence of (43) leads to the result that

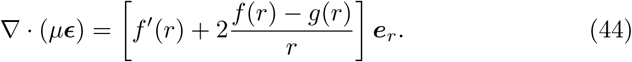

Further taking the curl of (44) yields equation (38). This completes our verification.

Note that equation (42) contains an arbitrary function, implying that the solution space of (38) is infinite-dimensional for this particular case. Therefore, it is impossible to uniquely determine *μ*, although the values of *μ* are known (*μ* = *μ*_0_) away from the cell surface. Accordingly, the solution for *λ* is *not* unique, since for an arbitrary *μ* given by (42), there exists an associated *λ* given by (39).

## Footnotes

1 In the area of biomechanics and mechanobiology, the techniques used to estimate cellular tractions have been referred to as “traction force microscopy (TFM).” We take exception to this nomenclature. We believe that “traction microscopy (TM)” alone is sufficient, and more importantly, accurate, since there is no “traction force” in mechanics. For this reason, in this manuscript we refer to “traction force microscopy” as “traction microscopy.”

